# Higher order assembling of the mycobacterial essential polar growth factor DivIVA/Wag31

**DOI:** 10.1101/394452

**Authors:** Komal Choukate, Aanchal Gupta, Brohmomoy Basu, Karman Virk, Munia Ganguli, Barnali Chaudhuri

## Abstract

How proteins localize at the pole remains an enigma. DivIVA/Wag31, which is an essential pole organizing protein in mycobacteria, can assemble at the negatively curved side of the membrane at the growing pole to form a higher order structural scaffold for maintaining cellular morphology and localizing various target proteins for cell-wall biogenesis. A single-site phosphorylation in Wag31 is linked to the regulation of peptidoglycan biosynthesis for optimal mycobacterial growth. The structural organization of polar scaffold formed by coiled-coil rich Wag31, which is a target for anti-tubercular agent amino-pyrimidine sulfonamide, remains to be determined. Here, we report biophysical characterizations of a phospho-mimetic (T73E) and a phospho-ablative (T73A) form of mycobacterial Wag31 using circular dichroism, small angle solution X-ray scattering and atomic force microscopy. While data obtained from both variants of Wag31 in solution states suggested formation of alpha-helical, large, elongated particles, their structural organizations were different. Atomic force microscopic images of Wag31 indicate polymer formation, with occasional curving, sharp bending and branching. Observed structural features in this first view of the polymeric forms of Wag31 suggest a basis for higher order network scaffold formation for polar protein localization.

## Introduction

Cell poles are sites for many critical biosynthetic and regulatory activities, which are important for polar growth, chemo-taxis and the proper progression of the cell cycle (1-3). How bacteria localize proteins at the cell pole is little understood. A ‘diffusion-and-capture’ process is suggested as one of the general mechanisms of recruitment at the pole, which must harbor a ‘hub’ or a ‘landmark’ for capturing the diffusing target entities (1-3). Examples of such pole-organizing ‘hub’ proteins include PopZ in *Caulobacter crescentus* and HubP in *Vibrio cholerae* (4-5).

Bacterial tropomyosin-like protein DivIVA recently emerged as one such important ‘hub’ or ‘landmark’, which is implicated in the polar capturing of a number of proteins and the chromosomal origin in several bacteria (6-10). DivIVA can sense curved, concave surfaces and can assemble at the inner, cytoplasmic side of the membrane at the pole or at the division septum to form a structural scaffold (7, 11-15). These structural scaffolds formed by DivIVA likely aid in the capturing and localization of various target proteins at the pole.

How do the ‘hub’ proteins like DivIVA sense concave membrane curvature and assemble at the pole? Proposed models range from factors such as geometric cues, physical or chemical factors, including difference in lipid composition, individual oligomerization as a curvature-sensing mechanism, collective interactions for curvature sensing, and volume exclusion (12, 16-18). It has been suggested that DivIVA localize at the pole by sensing geometric cue and utilizing higher order assembling (7, 12, 14, 19). However, little is known about these higher order oligomers of DivIVA.

Although curvature-sensitivity and coiled-coil organizations are common amongst DivIVAs, their biological roles can be very different in different bacteria (20). In *Bacillus subtilis*, DivIVA (bsDivIVA) participates in the localization of ‘Min’ division inhibitors (11). During sporulation in *B. subtilis*, bsDivIVA participates in polar anchoring of the chromosome origin, which is mediated by a DNA binding protein RacA (6, 21). In *Listeria monocytogenes*, DivIVA play multiple roles in secretion of autolysins, division septum positioning and swarming motility (22-23). In actinobacteria, DivIVA reportedly participates in cell-shape maintenance, origin tethering, protection against oxidative stress and directing the cell-wall biosynthetic proteins at the pole (15, 24-27). In *Corynebacterium glutamicum*, DivIVA localizes RodA at the pole (28). However, detailed natures of these protein-protein interactions are not known.

Human pathogen mycobacteria, unlike *Escherichia coli* and *B. subtilis*, grows exclusively at the pole, where the ‘old’ pole grows faster than the ‘new’ pole (29-30). DivIVA, which is also known as Wag31, is essential in mycobacteria and is required for the maintenance of cell morphology and for polar growth (15, 25, 31-32). Wag31 appears to play a critical role in polar growth by creating an excluded, inert zone at the pole and directing the localizations of some cytosolic cell-wall biogenesis enzymes at the sub-polar regions while interacting with others in mycobacteria (15). A single site (T73) phosphorylation of Wag31 relates to its better polar localization and higher peptidoglycan synthesis during rapid growth (32). At the pole, Wag31 acts as a molecular scaffold and interacts with many proteins involved in the cell wall synthesis and cell division such as acyl-CoA carboxylase, penicillin binding protein FtsI (a component of divisome), CwsA, and chromosomal origin partition protein ParA (15, 33-36). CwsA appears to play a role in Wag31 localization in mycobacteria (34). Furthermore, Wag31 is identified as a target for anti-tuberculosis agent amino-pyrimidine sulfonamide, with an unknown mechanism of action (37-38).

Structural information on DivIVA scaffolds is so far limited to only one protein from *B. subtilis*, presumbly due to the challenges in crystallizing a highly aggregation-prone, scaffold-forming protein. A tentative model of ∼30 nm long, curved, tetrameric bsDivIVA was obtained by combining an atomic-resolution structure of the N-terminal, lipid binding domain and a low-resolution structure of the C-terminal tetrameric domain containing only C-alpha atoms (14). Based on cryo-negative electron microscopy, a ∼22 nm long and ∼3 nm wide “doggy bone” shaped bsDivIVA structural unit was reported that further assembles to form higher order strings and network-like structures by lateral interaction (39). These higher order networks likely recruit target proteins at the pole and are important for morphology (39).

Here, we report first-ever *in vitro* characterizations of the phospho-mimetic and phospho-ablative forms of DivIVA/Wag31 from *M. tuberculosis* (tbWag31, Rv2145c from *M. tuberculosis*) using circular dichroism spectro-polarimetry (CD), atomic force microscopy (AFM) and small angle X-ray scattering (SAXS). While SAXS data are consistent with the formation of rod-shaped, elongated tbWag31 polymers, AFM images indicate that tbWag31 form polymeric structures with instances of bending and branching. We are presenting the first views of polymeric structures formed by tbWag31, that suggest how it can likely form higher order structural scaffold at the pole.

### Wag31 is alpha helical in solution

In order to learn about phosphorylation-dependent structural organizations of tbWag31, we purified full-length tbWag31 (∼28 kDa/monomer) in two forms, phospho-mimetic tbWag31-T73E and phospho-ablative tbWag31-T73A and confirmed by mass spectrometry (S1, supplementray material; 32, 40-41). To obtain the secondary structure contents of tbWag31 variants, we performed CD experiments (figure 1a). The CD profiles suggested high alpha-helical contents for both mutants, as expected for coiled-coil rich proteins. The two CD profiles of the mutants were slightly dissimilar (figure 1a), suggesting differences either at the level of secondary structure or in their respective assembly states caused by a single site mutation.

**Figure 1.**
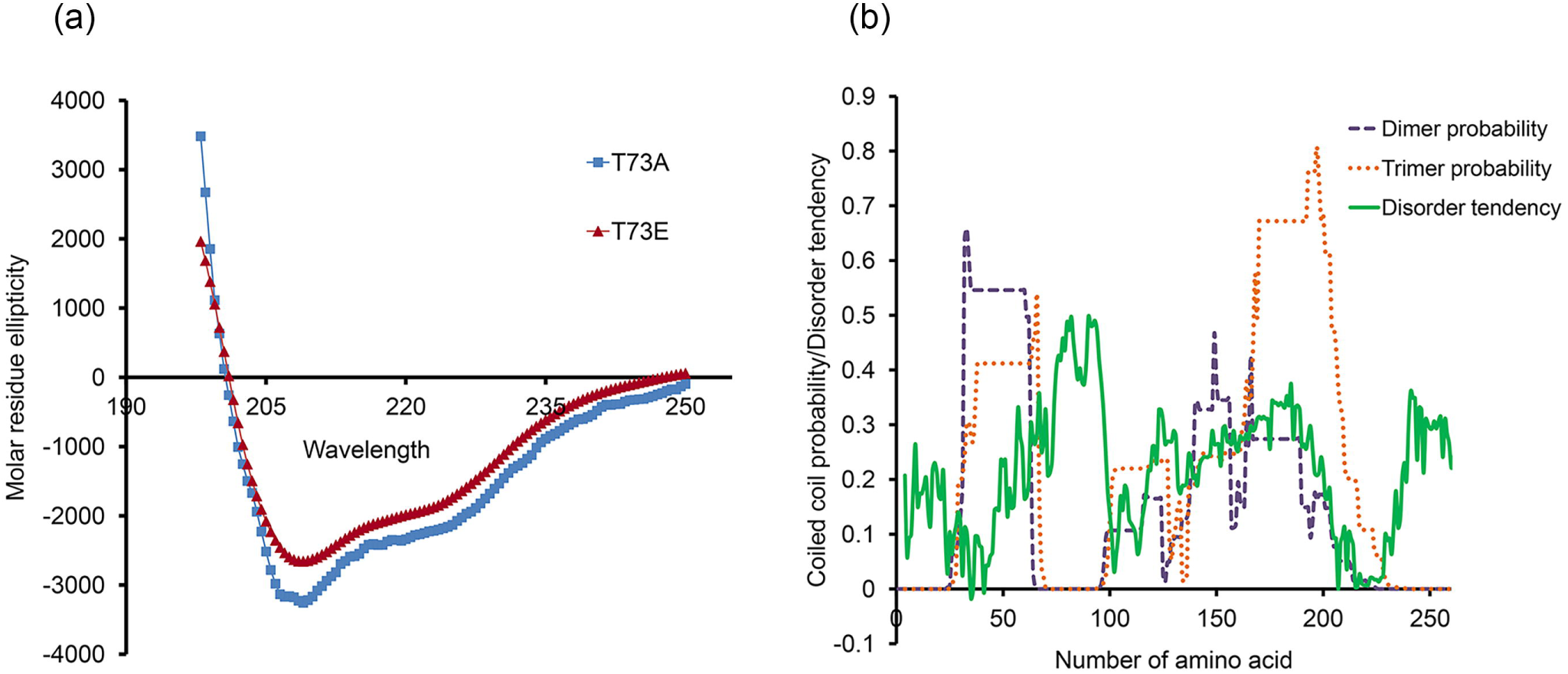
(a) Far-UV CD profiles of Wag31-T73E (red, triangle) and Wag31-T73A (blue, square) are shown (molar residue ellipticity in deg.cm^2^/dmol *versus* wavelength in nm). CD spectra (195-250 nm) of freshly purified proteins at 0.5 mg/ml concentrations dialyzed in 50 mM sodium phosphate buffer (pH 7.5) were recorded at 25° C on a Jasco J815 spectro-polarimeter in a cuvette with 0.1 cm pathlength. Three scans were averaged and analyzed by Dichroweb program (48). Due to the polymeric nature of our sample, estimations of alpha-helical contents were not performed. (b) Predicted coiled coil and intrinsic disordered regions *versus* tbWag31 sequence numbers are shown. Multicoil and IUPRED were used for coiled coil and intrinsic disorder predictions, respectively (49-50). Predicted disorder tendency values are shifted along the y-axis for better visualization.

Coiled coils can be assembled in many different ways, in parallel or anti-parallel orientations, to form dimer, trimer, tetramer, pentamer or hexamer (42), which can be easily exploited for higher order scaffold formation. Wag31-homolog bsDivIVA contains two coiled coil domains, a dimeric, N-terminal domain and a central, tetrameric C-terminal domain, that are joined by a short, ∼ 20 residue linker region to form a tetramer (14). A short, intertwined loop region in the terminally located N-terminal domain of tetrameric bsDivIVA interacts with the membrane (14). While a similar overall organization might be shared by these two proteins with ∼ 21 % sequence identity, tbWag31 (260 residues) is much longer than bsDivIVA (164 residues), and contains a phosphorylation target site (T73, 32). Two coiled coil domains were predicted for tbWag31: one in the N-terminal region and one in the C-terminal region, the latter showing higher propensity to be a trimer than a dimer (figure 1b, 31). The phosphorylated Thr73 residue in tbWag31 is housed within the ∼100 residue long linker region in between the two domains, which is predicted to be intrinsically disordered (figure 1b).

### Wag31 form elongated, rod-shaped structures in solution

Static SAXS data acquired from purified, 1-day old tbWag31-T73E and tbWag31-T73A proteins were analyzed to obtain model-free, averaged sizes of the scattering particles in solution (table 1, figure 2). Due to the polymeric nature of the tbWag31 proteins, routine SAXS data analysis applicable to globular proteins was not performed. Shapes of the modified Guinier plots for rods and dimensionless Kratky plots of the SAXS data were consistent with the presence of non-globular, elongated particles for both mutants (43-44). Cross-sectional radii of gyration (R_XS_) and linear mass densities (mass *per* unit length) were obtained from the slopes and intercepts of the modified Guinier plots (table 1; 44). It appears that the ablative mutant is much thinner than its mimetic counterpart in the cross-sectional direction. Possible reasons for this size difference could be lateral interactions between individual polymers or their conformational differences or branching or combinations thereof. On the other hand, linear mass densities of the polymeric mutants were not substantially different, thus ruling out lateral interaction (table 1, figure 2). The phospho-ablative mutant appears to be more rod-like than the phospho-mimetic mutant in the Kratky plot, which could be due to higher degree of branching in the latter (figure 2).

**Table 1.** Experimental conditions and results obtained from (a) SAXS and (b) AFM data analyses of Wag31 filaments. All concentrations were determined using Bradford assay. [NC: not calculated; SD: standard deviation; N: number of data points; ‘*same*‘: same conditions were used for mimetic and ablative mutants]

| (a) SAXS data analysis | tbWag31-T73E | tbWag31-T73A |
| --- | --- | --- |
| **Data-collection** |  |  |
| Experimental station | BM29, European Synchrotron Radiation facility, France (56) | *Same* |
| Wavelength (Å) | 1.0 | *Same* |
| Detector distance (m) | 2.9 | *Same* |
| Exposure time (sec) | 1 (up to 10 acquisitions) | *Same* |
| Concentration range (mg ml<sup>-1</sup>) | 2.7 | *Same* |
| Polymerization temperature (° C) | 4 | *Same* |
| Buffer | 50 mM Tris.HCl at pH 7.5, 300 mM NaCl, 10 % glycerol, 1mM EDTA and 5 mM BME | *Same* |
| Experimental temperature (° C) | 4 | *Same* |
| Software used | Primus and GNOM from the ATSAS suite (52) | *Same* |
| **Structural parameters**<br>(Calculated using Primus, ATSAS suite, 52) |  |  |
| R<sub>g</sub> (nm) [from Guinier plot, qR<sub>g</sub> < 1.3] | 19.8 | 21.8 |
| R<sub>xs</sub> (nm) [from modified Guinier plot for rods]. | 5.7 | 3.0 |
| Linear mass density (kDa/nm) [from modified Guinier plot for rods] | 13.1 | 12.6 |
| R<sub>xs</sub> (nm) [from P<sub>xs</sub>(r)] | 5.9 | 3.3 |
| Linear mass density (kDa/nm) | 13.0 | 12.8 |
| [from $P_{XS}(r)$] | | |
| $D_{XS-max}$ (nm) | 23.4 | 13.55 |
| **(b) AFM data analysis** | **tbWag31-T73E** | **tbWag31-T73A** |
| Instrument | 5500 scanning probe microscope (Keysight Technologies) at CSIR-IGIB, New Delhi | *Same* |
| Experimental parameters | Tapping mode in the air with 125-μm-long silicon cantilevers (AppNano);<br>Tip size 10 nm<br>Resonance frequency of 200-400 kHz;<br>Force constant of 13-77 Newton/m.<br>Scan speed: 1 line/s. | *Same* |
| Concentration range | 0.8 mg/ml stock solution (4° C) serially diluted to 0.004 mg/ml prior to the experiment | *Same* |
| Buffer | 50 mM Tris.HCl pH 7.5, 300 mM NaCl, 10% glycerol, 1 mM EDTA | *Same* |
| Software used | Keysight Technologies PicoView 1.20.3 software and PicoImage Elements software v7.3 | *Same* |
| **Structural parameters** |  |  |
| Average width (nm) | 45.0 (SD 16.0; N=25) | 52.0 (SD 13.0; N= 13) |
| Average height (nm) | 2.1 (SD 1.8; N= 72) | 1.4 (SD 0.9; N=17) |
| Angular range for Y-shaped branches (smallest angle) | 68° (18°; N=12)<br>Range: 50°-99° | NC |
| Angular range for sharp bents (smaller angle) | 97.8° (21°; N=10)<br>Range: 75°-146° | NC |

**Figure 2.**
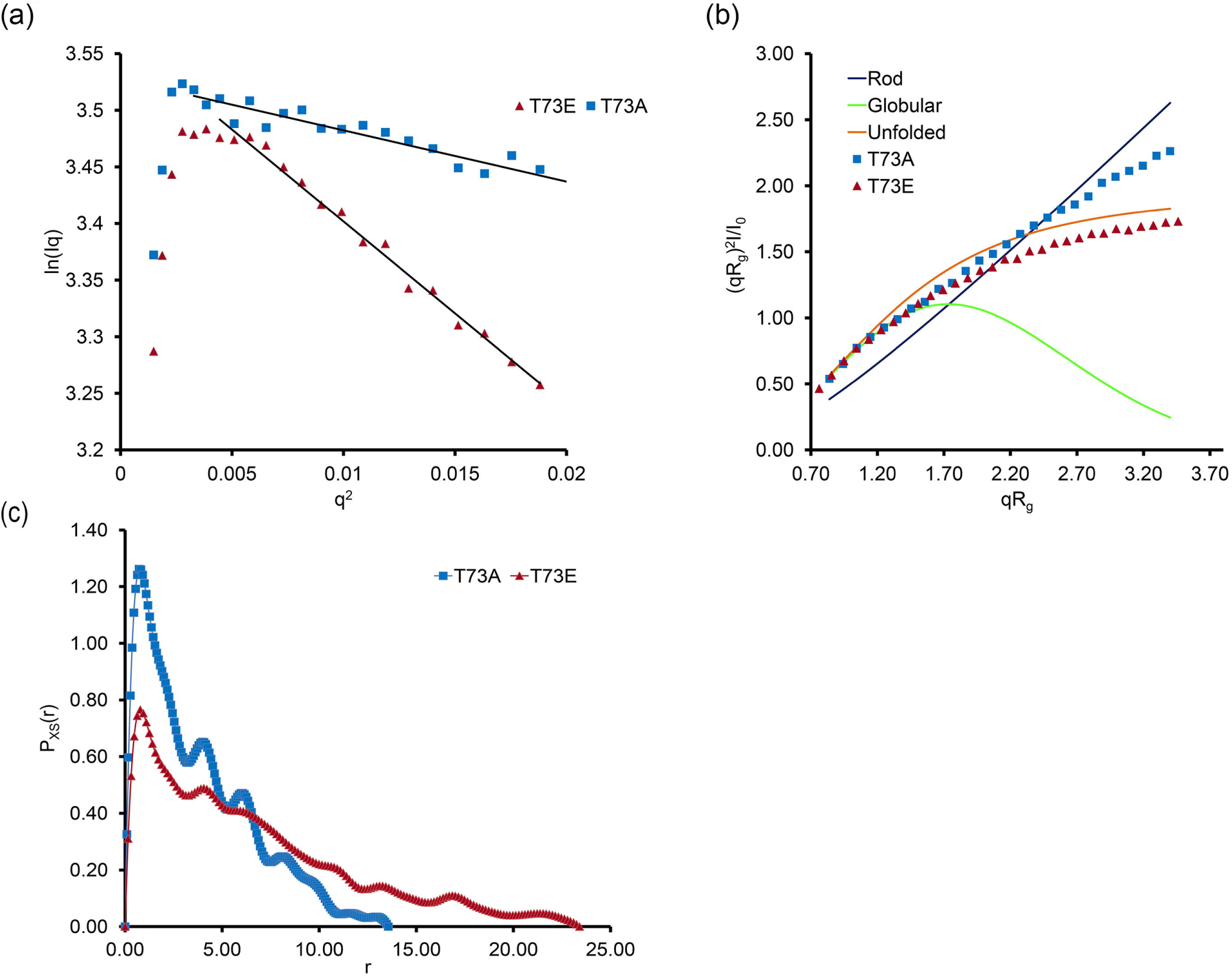
SAXS data of tbWag31-T73E (red, triangle) and tbWag31-T73A (blue, square). (a) Modified Guinier plots for rod-shaped particles (LnI(q).q *versus* q^2^) are shown, where I is the intensity and q is the momentum transfer in nm^−1^ (q= 4.π.sin(θ)/λ, 2θ is the scattering angle, λ is the X-ray wavelength). (b) Dimensionless Kratky plots (I(q)/I(0).(q.R_g_)^2^ *versus* (q.R_g_), where I(0) is the forward scattering and R_g_ is the radius of gyration as obtained from the Guinier plots (q.R_g_ < 1.3). Theoretical Kratky plots for globular, coil and rod-shaped scattering particles are shown for comparison (51). (c) Cross-sectional pair-distribution function (P_XS_) *versus* r in nm (option 4 in GNOM from the ATSAS suite, 52), where r is the pair-wise distance in real space. P_XS_ functions were calculated under boundary conditions that P_XS_=0 at r=0 and at r ≥ D_XS-max_, D_XS-max_ being the maximum particle diameter in the cross-sectional direction. SAXS datasets are deposited in Small Angle Scattering Biological Databank (SASBDB), with accession numbers SASDEB2 and SASDEC2 (53).

A comparison of cross-sectional pair distribution functions (P_XS_) further supported difference in cross-sectional sizes between the tbWag31-T73E and tbWag31-T73A mutants (table 1, figure 2). Cross-sectional pair distribution functions suggest a substantial change in structural organization between the two mutants, with a large (∼ 9.8 nm) difference in their maximum cross-sectional diameters. Interestingly, the first two peaks in both the P_XS_ functions are co-located (∼ 0.6 nm and 3.8 nm). It is possible that the phospho-mimetic mutant has more extended conformation and/or more branched than the ablative form, which is consistent with the differences observed in their modified Guinier, P_XS_ and Kratky plots (figure 2). All the solution size parameters are summarized in table 1a.

### Wag31 assemble to form polymers with bending and branching

AFM provided a wealth of information on coiled-coil rich intermediate filaments (45-46). To directly visualize the higher order forms of Wag31, AFM imaging of 2-7 days old tbWag31 samples were performed (table 1, figure 3). Protein samples were serially diluted in water to a final concentration of 0.004 mg/ml, applied to a freshly cleaved mica surface and air dried prior to imaging (table 1). Filaments were observed for both variants of tbWag31 (figure 3). However, noticeably fewer tbWag31-T73A filaments were located on the mica surface than tbWag31-T73E filaments under comparable experimental conditions. Therefore, most of our discussions will be relevant to the phospho-mimetic form of tbWag31. Lengths of the polymers changed substantially between 2^nd^ and 5^th^ day old tbWag31-T73E (more than 10 micron), indicating primarily linear growth. Fewer long polymers were detected on 6^th^ and 7^th^ day.

**Figure 3.**
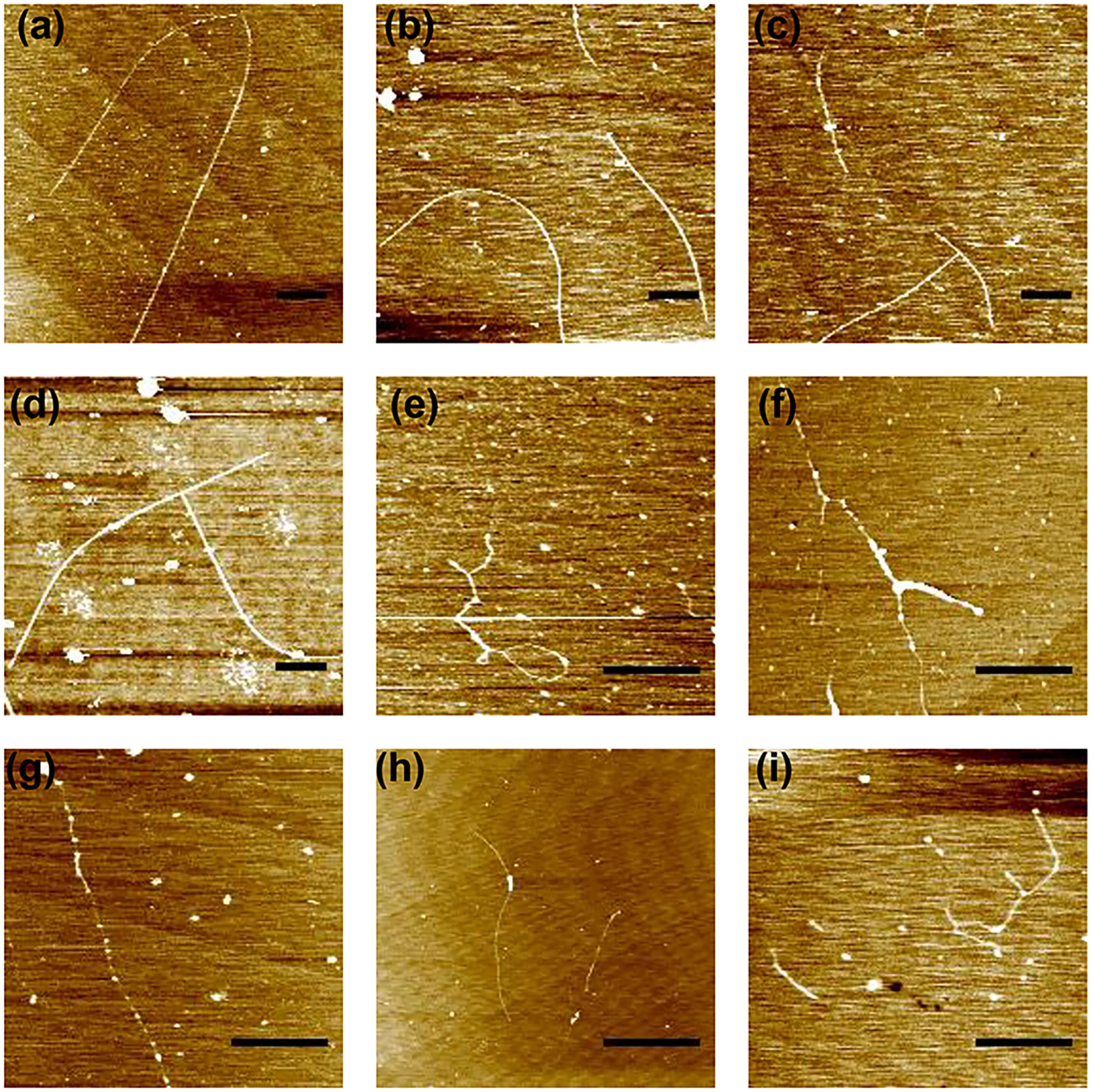
AFM topographic images of (a-f) tbWag31-T73E and (g-i) tbWag31-T73A filaments. Minimum image processing (first order flattening and brightness contrast) was employed. Contour lengths were measured by tracing the molecular contour with a segmented line. All heights were measured in nm as the difference between the top of the biomolecules and the average height of the underlying mica using PicoImage Elements software v7. We note that AFM images have lower resolution in the horizontal direction (width, because of the limit on resolution due to the tip width) than the vertical (height) direction, typically causing apparent width broadening (54). In addition, measured heights are influenced by electrostatic interactions (55). Scale bars are one micron.

Most interesting features of both the tbWag31 filaments were occurrences of occasional curving, kinking (sharp change in filament direction) and apparently branched structures (figure 3). These bent/branched structures can be formed by dimeric, trimeric or other forms of coiled coils. Measured bending (smaller) and branching (smallest) angles in the tbWag31-T73E filaments showed wide-ranging distributions (table 1).

Measured heights obtained from the AFM images show rather wide cross-sectional size range for the phospho-mimetic form of tbWag31 (table 1). No clear temporal pattern in height changes were detected for the tbWag31-T73E filaments obtained from 2^nd^ to 7^th^ day. Due to fewer instances of tbWag31-T73A polymers in comparison to the tbWag31-T73E polymers in the acquired AFM images, we cannot comment on any difference in height or the level of branching/bending between these two polymers. Results of AFM image analysis are provided in table 1.

## Summary

We present low resolution structural characterizations of higher order assemblies formed by the phospho-mimetic and phospho-ablative variants of tbWag31, based on CD, SAXS and AFM data. While SAXS data provided estimations of averaged cross-sectional sizes of hydrated tbWag31 polymers in solution, AFM images allowed direct visualizations of air-dried polymers on the mica surface. As the AFM images were acquired at much lower concentrations of proteins than that used for the SAXS experiments, we are not performing any direct comparison of results derived from these experiments. The phosphor-ablative form appears to have different structural organization (SAXS) and less frequently seen (AFM) than the phosphor-mimetic form of the polymer. Observed differences between the two mutants are in accordance with previously published yeast two-hybrid data that the phospho-mimetic Wag31 is relatively better at self-interaction amongst the two, that likely relates to phosphorylation-dependent regulation of cell wall synthesis (32).

The AFM images presented here suggest how a Wag31 scaffold might form at the cell pole. It has been reported that bsDivIVA forms bent tetrameric structure that makes rather tenuous contact with the membrane (14). In addition, bsDivIVA can form “doggy-bone” shaped structural units with bifurcated ends and can form 2-dimensional structures by lateral interaction of strings composed of “doggy-bone” units (39). An ability to form curved polymer with a curvature perfectly matching that of the polar curvature of the cell was suggested as one of the possible mechanisms of polar assembling (39). Although we observed polymers that appear to be curved and branched, our data shows no clear evidence of lateral interaction between these polymers. Considering the diameter of rod-shaped mycobacteria (of the order of 300-500 nm, 47), the several micron long extended filaments observed *in vitro* are probably results of unimpeded filament growth under experimental conditions. Nevertheless, these AFM images provided very important clues about possible Wag31-mediated scaffold formation at the pole. These tbWag31 polymers appear to be nimble and capable of curving, kinking and branching with a wide range of angles (table 1, figure 3). Recurring occurrences of the observed structural features in a growing Wag31 polymer can likely lead to the formation of a localized networked structure within the polar confinement (figure 4). Atomic details of these structures, effect of phosphorylation and lipid bilayer on these structures, the frequency and structural determinants of bending/branching with respect to the polar membrane and directionality of polymer growth, remains to be explored.

**Figure 4.**
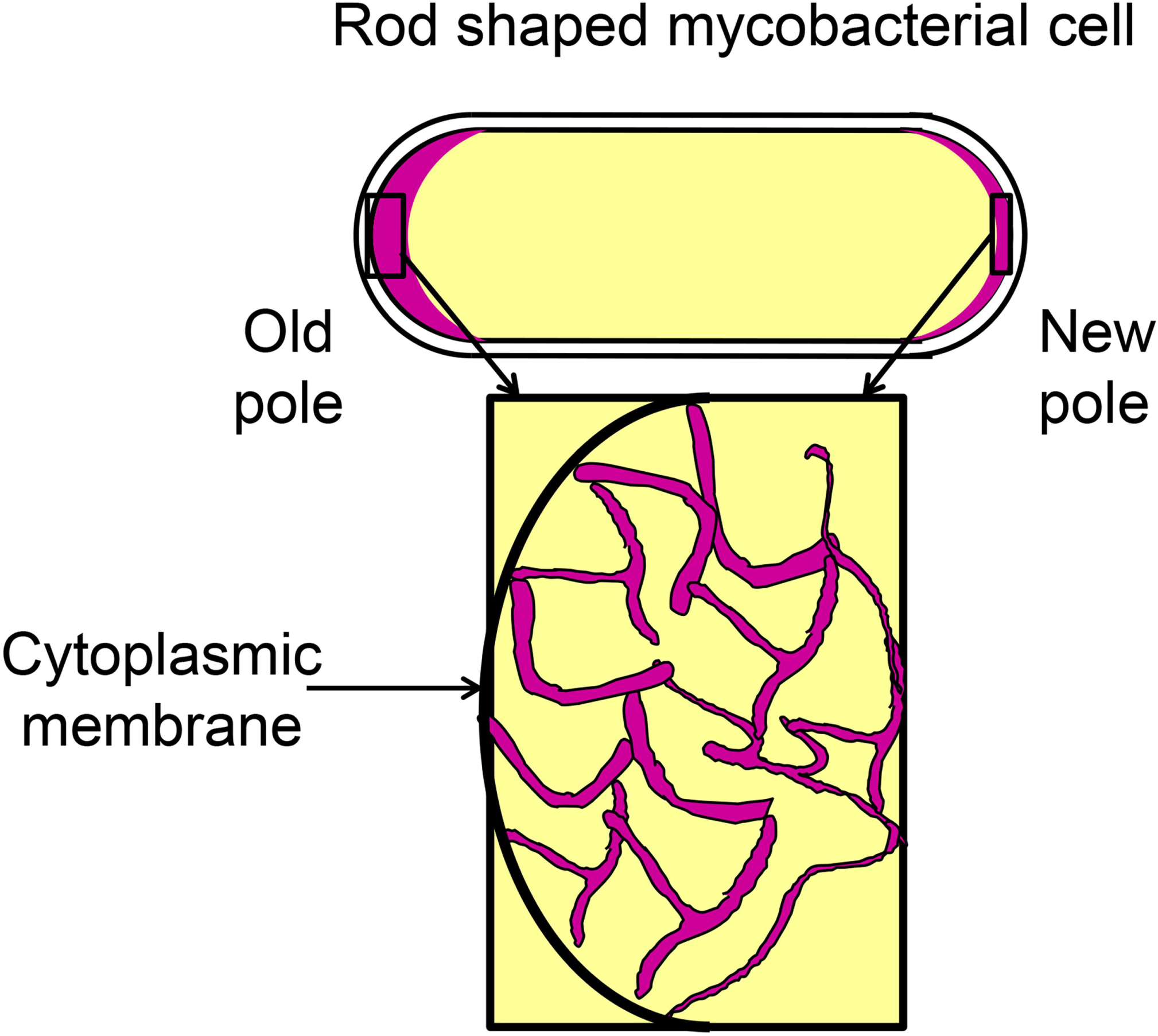
A cartoon diagram showing probable polar Wag31 scaffold (in maroon) formed by a filament with curving, bending and branching.

## Acknowledgements

Authors acknowledge intramural CSIR support. KC and AG are recipients of CSIR and DBT research fellowships, respectively. Access to SAXS data collection facility at the BM29 beamline, European Synchrotron Radiation Facility (ESRF) in France was supported by Department of Biotechnology (Government of India).

## References

1. Shapiro, L., McAdams, H. H. & Losick, R. (2009). Why and how bacteria localize proteins. Science. 326, 1225–1228.

2. Rudner, D. Z. & Losick, R. (2010). Protein Subcellular Localization in Bacteria. Cold Spring Harb Perspect Biol. 2(4): a000307.

3. Laloux, G. & Jacobs-Wagner, C. (2014). How do bacteria localize proteins to the cell pole? J. Cell Sci. 127, 11–19.

4. Bowman, G. R., Comolli, L. R., Zhu, J., Eckart, M., Koenig, M., Downing, K. H., Moerner, W. E., Earnest, T. & Shapiro, L. (2008). A polymeric protein anchors the chromosomal origin/ParB complex at a bacterial cell pole. Cell 134, 945–955.

5. Yamaichi, Y., Bruckner, R., Ringgaard, S., Möll, A., Cameron, D. E., Briegel, A., Jensen, G. J., Davis, B. M. & Waldor, M. K. (2012). A multidomain hub anchors the chromosome segregation and chemotactic machinery to the bacterial pole. Genes Dev. 26, 2348–2360.

6. Ben-yehuda, S., Rudner, D. Z. & Losick, R. (2003). RacA, a bacterial protein that anchors chromosomes to the cell poles. Science. 299, 532–536.

7. Lenarcic, R., Halbedel, S., Visser, L., Shaw, M., Wu, L. J., Errington, J., Marenduzzo, D. & Hamoen, L. W. (2009). Localisation of DivIVA by targeting to negatively curved membranes. EMBO J. 28, 2272–2282.

8. Letek, M., Ordóñez, E., Vaquera, J., Margolin, W., Flärdh, K., Mateos, L. M. & Gil, J. A. (2008). DivIVA is required for polar growth in the MreB-lacking rod-shaped actinomycete Corynebacterium glutamicum. J. Bacteriol. 190, 3283–3292.

9. Letek, M., Fiuza, M., Villadangos, A. F., Mateos, L. M. & Gil, J. A. (2012). Cytoskeletal proteins of actinobacteria. Int. J. Cell Biol. 2012.

10. Lin, L. & Thanbichler, M. (2013). Nucleotide-independent cytoskeletal scaffolds in bacteria. Cytoskeleton 70, 409–423.

11. Edwards, D. H. & Errington, J. (1997). The Bacillus subtilis DivIVA protein targets to the division septum and controls the site specificity of cell division. Mol. Microbiol. 24, 905–915.

12. Ramamurthi, K. S. & Losick, R. (2009). Negative membrane curvature as a cue for subcellular localization of a bacterial protein. Proc. Natl. Acad. Sci. 106, 13541–13545.

13. Ramamurthi, K. S. (2010). Protein localization by recognition of membrane curvature. Curr Opin Microbiol. 13, 753–7.

14. Oliva, M. A., Halbedel, S., Freund, S. M., Dutow, P., Leonard, T. A., Veprintsev, D. B., Hamoen, L. W. & Löwe, J. (2010). Features critical for membrane binding revealed by DivIVA crystal structure. EMBO J. 29, 1988–2001.

15. Meniche, X., Otten, R., Siegrist, M. S., Baer, C. E., Murphy, K. C., Bertozzi, C. R. & Sassetti, C. M. (2014). Subpolar addition of new cell wall is directed by DivIVA in mycobacteria. Proc. Natl. Acad. Sci. 111, E3243–E3251.

16. Mukhopadhyay, R., Huang, K.C., and Wingreen, N.S. (2008). Lipid localization in bacterial cells through curvature-mediated microphase separation. Biophys. J. 95, 1034–1049.

17. Strahl, H., Ronneau, S., Gonzalez, B.S., Klutsch, D., Schaffner-Barbero, C., & Hamoen, L.W. (2015). Transmembrane protein sorting driven by membrane curvature. Nat. Commun. 6, 8728.

18. Surovtsev, I. V. & Jacobs-Wagner, C. (2018). Subcellular Organization: A Critical Feature of Bacterial Cell Replication. Cell. 172, 1271–1293.

19. Wagstaff, J. & Löwe, J. (2018). Prokaryotic cytoskeletons: Protein filaments organizing small cells. Nat. Rev. Microbiol. 16, 187–201.

20. Kaval, K. G. & Halbedel, S. (2012). Architecturally the same, but playing a different game: the diverse species-specific roles of DivIVA proteins. Virulence 3, 406–407.

21. Wu, L. J. & Errington, J. (2003). RacA and the Soj-Spo0J system combine to effect polar chromosome segregation in sporulating Bacillus subtilis. Mol. Microbiol. 49, 1463– 1475.

22. Kaval, K. G., Rismondo, J. & Halbedel, S. (2014). A function of DivIVA in Listeria monocytogenes division site selection. Mol. Microbiol. 94, 637–654.

23. Kaval, K. G., Hauf, S., Rismondo, J., Hahn, B. & Halbedel, S. (2017). Genetic dissection of DivIVA functions in Listeria monocytogenes. J. Bacteriol. 199, e00421–17.

24. Hempel, A. M., Wang, S. B., Letek, M., Gil, J. A. & Flärdh, K. (2008). Assemblies of DivIVA mark sites for hyphal branching and can establish new zones of cell wall growth in Streptomyces coelicolor. J Bacteriol. 190 (22), 7579–83.

25. Kang, C. M., Nyayapathy, S., Lee, J. Y., Suh, J. W. & Husson, R. N. (2008). Wag31, a homologue of the cell division protein DivIVA, regulates growth, morphology and polar cell wall synthesis in mycobacteria. Microbiology 154, 725–735.

26. Mukherjee, P., Sureka, K., Datta, P., Hossain, T., Barik, S., Das, K. P., Kundu, M. & Basu, J. (2009). Novel role of Wag31 in protection of mycobacteria under oxidative stress. Mol. Microbiol. 73, 103–119.

27. Donovan, C., Sieger, B., Krämer, R. & Bramkamp, M. (2012). A synthetic Escherichia coli system identifies a conserved origin tethering factor in Actinobacteria. Mol. Microbiol. 84, 105–116.

28. Sieger, B., Schubert, K., Donovan, C. & Bramkamp, M. (2013). The lipid II flippase RodA determines morphology and growth in Corynebacterium glutamicum. Mol. Microbiol. 90, 966–982.

29. Kieser, K. J. & Rubin, E. J. (2014). How sisters grow apart: Mycobacterial growth and division. Nat. Rev. Microbiol. 12, 550–562.

30. Cameron, T. A., Zupan, J. R. & Zambryski, P. C. (2015). The essential features and modes of bacterial polar growth. Trends Microbiol. 23, 347–353.

31. Nguyen, L., Scherr, N., Gatfield, J., Walburger, A., Pieters, J. & Thompson, C. J. (2007). Antigen 84, an effector of pleiomorphism in Mycobactenum smegmatis. J. Bacteriol. 189, 7896–7910.

32. Jani C., Eoh H., Lee J. J., Hamasha K., Sahana M. B., Han J. S., Nyayapathy S., Lee J. Y., Suh J. W., Lee S. H., Rehse S. J., Crick D. C., Kang C. M. (2010). Regulation of polar peptidoglycan biosynthesis by Wag31 phosphorylation in mycobacteria. BMC Microbiol. 10.

33. Mukherjee, P., Sureka, K., Datta, P., Hossain, T., Barik, S., Das, K. P., Kundu, M., Basu, J. (2009). Novel role of Wag31 in protection of mycobacteria under oxidative stress. Mol Microbiol. 73(1), 103–19.

34. Plocinski P., Arora N., Sarva K., Blaszczyk E., Qin H., Das N., Plocinska R., Ziolkiewicz M., Dziadek J., Kiran M., Gorla P., Cross T. A., Madiraju M., Rajagopalan M.(2012). Mycobacterium tuberculosis CwsA interacts with CrgA and Wag31, and the CrgA-CwsA complex is involved in peptidoglycan synthesis and cell shape determination. J. Bacteriol. 194, 6398–6409.

35. Ginda, K., Bezulska, M., Ziólkiewicz, M., Dziadek, J., Zakrzewska-Czerwińska, J. & Jakimowicz, D. (2013). ParA of Mycobacterium smegmatis co-ordinates chromosome segregation with the cell cycle and interacts with the polar growth determinant DivIVA. Mol. Microbiol. 87, 998–1012.

36. Xu, W. X., Zhang, L., Mai, J. T., Peng, R. C., Yang, E. Z., Peng, C. & Wang, H. H. (2014). The Wag31 protein interacts with AccA3 and coordinates cell wall lipid permeability and lipophilic drug resistance in Mycobacterium smegmatis. Biochem. Biophys. Res. Commun. 448, 255–260.

37. Singh V., Dhar N., Pató J., Kolly G. S., Korduláková J., Forbak M., Evans J. C., Székely R., Rybniker J., Palčeková Z., Zemanová J., Santi I., Signorino-Gelo F., Rodrigues L., Vocat A., Covarrubias A. S., Rengifo M. G., Johnsson K., Mowbray S., Buechler J., Delorme V., Brodin P., Knott G. W., Aínsa J. A., Warner D. F., Kéri G., Mikušová K., McKinney J. D., Cole S. T., Mizrahi V., Hartkoorn R. C. (2017). Identification of aminopyrimidine-sulfonamides as potent modulators of Wag31-mediated cell elongation in mycobacteria. Mol. Microbiol. 103, 13–25.

38. Boshoff, H. I. (2017). Caught between two proteins: a mycobacterial inhibitor challenges the mold. Mol. Microbiol. 103, 2–6.

39. Stahlberg, H., Kutejová, E., Muchová, K., Gregorini, M., Lustig, A., Müller, S. A., Olivieri, V., Engel, A., Wilkinson, A. J. & Barák, I. (2004). Oligomeric structure of the Bacillus subtilis cell division protien DivIVA determined by transmission microscopy. Mol. Microbiol. 52, 1281–1290.

40. Lee, J. J., Kan, C. M., Lee, J. H., Park, K. S., Jeon, J. H. & Lee, S. H. (2014). Phosphorylation-dependent interaction between a serine/threonine kinase PknA and a putative cell division protein Wag31 in Mycobacterium tuberculosis. New Microbiol. 37, 525–33.

41. Perkins, D. N., Pappin, D. J., Creasy, D. M. & Cottrell, J. S. (1999). Probability-based protein identification by searching sequence databases using mass spectrometry data. Electrophoresis. 20, 3551–3567

42. Moutevelis, E. & Woolfson, D. N., (2009). A periodic table of coiled-coil protein structures. J Mol Biol. 385(3), 726–32.

43. Durand, D., Vivès, C., Cannella, D., Pérez, J., Pebay-Peyroula, E., Vachette, P. & Fieschi, F. (2010). NADPH oxidase activator p67(phox) behaves in solution as a multidomain protein with semi-flexible linkers. J. Struct. Biol. 169, 45–53.

44. Qian, S., Dean, R., Urban, V. S., Chaudhuri, B. N. (2012). The internal organization of mycobacterial partition assembly: does the DNA wrap a protein core? PLoS One. 7, e52690.

45. Herrmann, H., Kreplak, L. & Aebi, U. (2004). Isolation, characterization, and in vitro assembly of intermediate filaments. Methods Cell Biol. 78, 3–24.

46. Brodehl, A., Dieding, M., Klauke, B., Dec E., Madaan, S., Huang, T., Gargus, J., Fatima, A., Saric, T., Cakar, H., Walhorn, V., Tönsing, K., Skrzipczyk, T., Cebulla, R., Gerdes, D., Schulz, U., Gummert, J., Svendsen, J. H., Olesen, M. S., Anselmetti, D., Christensen, A. H., Kimonis, V., Milting, H. (2013). The novel desmin mutant p.A120D impairs filament formation, prevents intercalated disk localization, and causes sudden cardiac death. Circ Cardiovasc Genet. 6 (6), 615–23.

47. Cook, G. M., Berney, M., Gebhard, S., Heinemann, M., Cox, R. A., Danilchanka, O., Niederweis, M. (2009). Physiology of mycobacteria. Adv Microb Physiol. 55, 81–182.

48. Whitmore, L. & Wallace, B. A. (2004). DICHROWEB, an online server for protein secondary structure analyses from circular dichroism spectroscopic data. Nucleic Acids Research. 32, W668–673.

49. Wolf, E., Kim, P. S. & Berger, B. (1997). MultiCoil: a program for predicting two-and three-stranded coiled coils. Protein Sci. 6, 1179–1189.

50. Dosztányi, Z., Csizmok, V., Tompa, P. & Simon, I. (2005). IUPred: Web server for the prediction of intrinsically unstructured regions of proteins based on estimated energy content. Bioinformatics. 21, 3433–3434.

51. Glatter, O. & Kratky, O., eds. (1982). Small-angle X-ray Scattering. Academic Press, London.

52. Franke, D., Petoukhov, M. V., Konarev, P. V., Panjkovich, A., Tuukkanen, A., Mertens, H. D. T., Kikhney, A. G., Hajizadeh, N. R., Franklin, J. M., Jeffries, C. M. & Svergun, D. I. (2017). ATSAS 2.8: a comprehensive data analysis suite for small-angle scattering from macromolecular solutions. J Appl Crystallogr. 50, 1212–1225.

53. Valentini, E., Kikhney, A. G., Previtali, G., Jeffries, C. M. & Svergun, D. I. (2015). SASBDB, a repository for biological small-angle scattering data. Nucleic Acids Res. 43, D357–363.

54. van Noort, S. J. T., van der Werf, K., de Grooth, B. G., van Hulst, N. F. & Greve, J (1997). Height anomalies in tapping mode atomic force microscopy in air caused by adhesion. Ultramicroscopy, 69 (2), 117–127.

55. Müller, D. J. & Engel, A., (1997). The height of biomolecules measured with the atomic force microscope depends on electrostatic interactions. Biophys J. 73(3), 1633–44.

56. Brennich, M., Pernot, P, & Round, A. (2017). How to Analyze and Present SAS Data for Publication. Adv Exp Med Biol. 1009, 47–64.

